# Multiscale analysis of 3D nuclear morphology reveals new insights into growth plate organization in mice

**DOI:** 10.1101/375949

**Authors:** Sarah Rubin, Tomer Stern, Paul Villoutreix, Johannes Stegmaier, Yoseph Addadi, Elazar Zelzer

## Abstract

The shape of the nucleus is tightly associated with cell morphology, the mechanical environment, and differentiation and transcriptional states. Yet, imaging of nuclei in three dimensions while preserving the spatial context of the tissue has been highly challenging. Here, using the embryonic tibial growth plate as a model for cell differentiation, we study nuclear morphology by imaging cleared samples by light-sheet fluorescence microscopy. Next, we quickly segmented tens of thousands of nuclei using several open-source tools including machine learning. Finally, segmented nuclei underwent morphometric analysis and 3D spatial reconstruction using newly designed algorithms. Our method revealed differences in nuclear morphology between chondrocytes at different differentiation stages. Additionally, we identified different morphological patterns in opposing growth plates, such as gradients of volume and surface area, as well as features characteristic of specific growth plate zones, such as sphericity and orientation. Altogether, this work supports a link between nuclear morphology and cell differentiation. Moreover, it demonstrates the suitability of our approach for studying the relationships between nuclear morphology and organ development.

**Author summary:** There has been a growing interest in the relationship between nuclear morphology and its regulation of gene expression. However, to study global patterns of nuclear morphology within a tissue we must address the problem of acquiring and analyzing multiscale data, ranging from the tissue level through to subcellular resolution. We have established a new pipeline that enables acquisition and segmentation of hundreds of thousands of nuclei at a resolution that allows quantitative analysis. Moreover we have developed new algorithms that allow superimposing morphological aspects of hundreds of thousands of nuclei onto a visual representation of the entire tissue, allowing us to study nuclear morphology at an organ level. Using mouse growth plates as a model for the relationship between nuclear morphology and tissue differentiation, we show that nuclei change different aspects of their morphology during chondrocyte differentiation. Growth plates are usually described generically in the literature, suggesting they lack unique characteristics. We challenge this dogma by showing that morphological features such as volume distribute differently in opposing growth plates. Altogether, this work highlights the possible role of nuclear shape in the regulation of cell differentiation and demonstrates that our approach enables the study of nuclear morphology patterns within a tissue.

## Introduction

The ability to analyze multiscale data is essential for investigating the contribution of cellular components to tissue morphogenesis. Analyzing three-dimensional (3D) data becomes complicated when they span multiple scales, as is the case of the spatial relationships between cells and tissue. Focusing on the tissue level results in loss of subcellular resolution, while focusing on smaller regions at high resolution results in loss of their relationship to 3D tissue morphology. To cover all these scales, the entire dataset must be at the highest desired resolution, which produces even more challenges regarding data size. To study the cellular contribution to tissue morphogenesis, we need a way to easily explore large datasets describing the tissue level, while preserving individual cell features.

To image whole tissues and visualize in 3D subcellular tissue components, optical sectioning is often applied. Using fluorescence microscopes such as confocal or two-photon, this method produces high spatial-resolution datasets, but at the cost of low imaging speed. Moreover, these techniques are limited in imaging depth and are most suitable for tissues up to 300 μm in thickness. Light-sheet fluorescence microscopy (LSFM), which quickly acquires large volumes of tissue at submicron scale, is ideal for imaging fluorescent tissues or even whole organisms such as zebrafish (1, 2), *C.elegans* (3, 4), or *Drosophila* (5). Whole organism imaging with LSFM is possible because these model organisms are transparent and small, a critical factor for this modality. Thus, applying LSFM to opaque mammalian tissues requires tissue clearing to enable in-depth imaging.

Biological samples absorb and scatter light. Tissue clearing is the process of reducing light scattering and absorption by making tissues optically transparent. This process, which typically involves removing macromolecules and matching refractive indices within samples, results in better penetration of light through the sample, thereby increasing imaging depths (6). Recently, several organs from adult mice were successfully cleared using aqueous-based buffers, allowing for imaging up to 1.5 mm in depth while maintaining endogenous fluorescence (7-9). Combining tissue clearing with LSFM will allow for high-resolution mapping of fine tissue architectures in an intact 3D sample (10).

For many years, researchers have used information about nuclei to extrapolate about the state of a tissue. Changes in nuclear morphology have been associated with malignant transformation of cells and tissue (11, 12), aging (13, 14), cell differentiation (15-18), and response to mechanical forces (19-21). In the last 15 years, accumulating data have connected nuclear morphology with molecular mechanisms that involve gene expression or transcription (19, 22-24). Chromatin is organized into open and closed domains, which allow for the regulation of gene expression (25). The finding that chromatin associates with nuclear lamins and with the inner nuclear membrane provides a mechanism whereby alterations in nuclear morphology can regulate gene expression (26). These findings highlight the importance of studying global patterns of nuclear morphology within a tissue.

The growth plate is an excellent model system for studying the multiscale relationships among nuclear morphology, cell differentiation and tissue geometry. Growth plates are cartilaginous tissues located at either end of developing bones and responsible for their elongation. Starting in embryonic development, this tissue comprises chondrocytes that display a conserved spatial organization into zones, namely the resting (RZ), proliferative (PZ), prehypertrophic (PHZ), and hypertrophic zones (HZ). These zones, reflecting a series of differentiation states, are marked by unique cell morphologies, extracellular matrix (ECM) properties and gene expression profiles (27-31).

Despite these advantages, composing a multiscale dataset from growth plates poses several challenges. Growth plates are extremely dense, as they are packed with collagen-rich extracellular matrix (31), which hinders 3D imaging. Additionally, the tissue is relatively large, reaching >1 mm in thickness, which implies longer durations of imaging and data analysis. Finally, current methodologies to segment nuclei are not suitable for large datasets, within which image quality may vary, resulting in slow and unreliable segmentations. In this work, we address these challenges using an especially designed pipeline for 3D imaging and analysis of growth plate cell nuclei as well as their spatial relationships within the tissue. We imaged nuclei from cleared embryonic growth plates using LSFM, applied several open-source tools to quickly segment nuclei, and developed a visualization platform to explore our obtained data. Our method revealed that during chondrocyte differentiation, nuclei underwent several morphological changes that were common to all growth plates, such as in nuclear orientation. By contrast, we identified different patterns in the two opposing growth plates in features such as nuclear volume and surface area.

## Results

### A pipeline for imaging the growth plate and quantifying large multiscale data

To study nuclear morphology in an intact mouse growth plate, we developed a high-throughput method that allowed us to characterize nuclear morphology quantitatively while maintaining the spatial relation of each nucleus within the tissue (Fig 1). To overcome the challenge of imaging a large and dense tissue like the growth plate, we cleared the tissue using the simple and inexpensive PACT-deCAL technique (7, 8), which was found to allow nuclear labeling while preserving tissue fluorescence. Next, embryonic day (E) 16.5 DAPI-labeled tibias were imaged by light-sheet microscopy (Fig 2A). Due to its high speed, light-sheet microscopy proved to be the superior choice over other laser-based techniques, such as confocal and two-photon microscopy. To ensure that light would pass through the sample, we used dual side illumination when necessary and rotated the sample to find the optimal imaging orientation. To reduce the file size and expedite image processing, acquired z stacks included only regions with growth plate nuclei.

**Figure 1.**
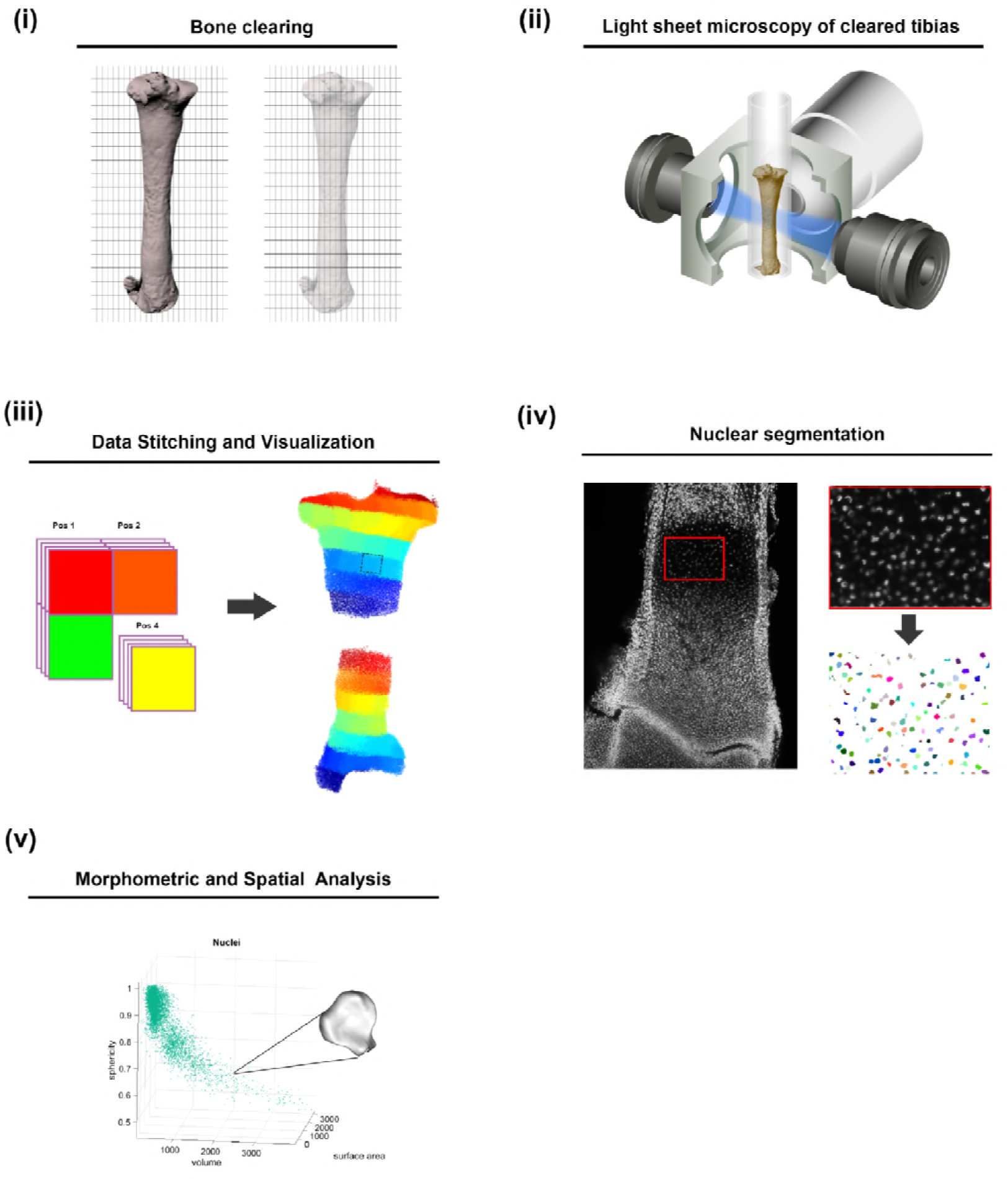
A pipeline for imaging growth plates and quantify large multiscale data in 3D. The bone is dissected and cleared with PACT-deCAL, and nuclei are fluorescently labeled. (ii) The cleared bone is embedded in a glass capillary and imaged at low resolution, followed by high-resolution imaging of the proximal and distal growth plates with a Z.1 light sheet microscope. (iii) Each image stack undergoes nuclear segmentation. (iv) The coordinates of each segmented nucleus are mapped back to their anatomical position in the bone and (v) the segmented nuclei are subjected to morphometric and spatial analyses.

**Figure 2.**
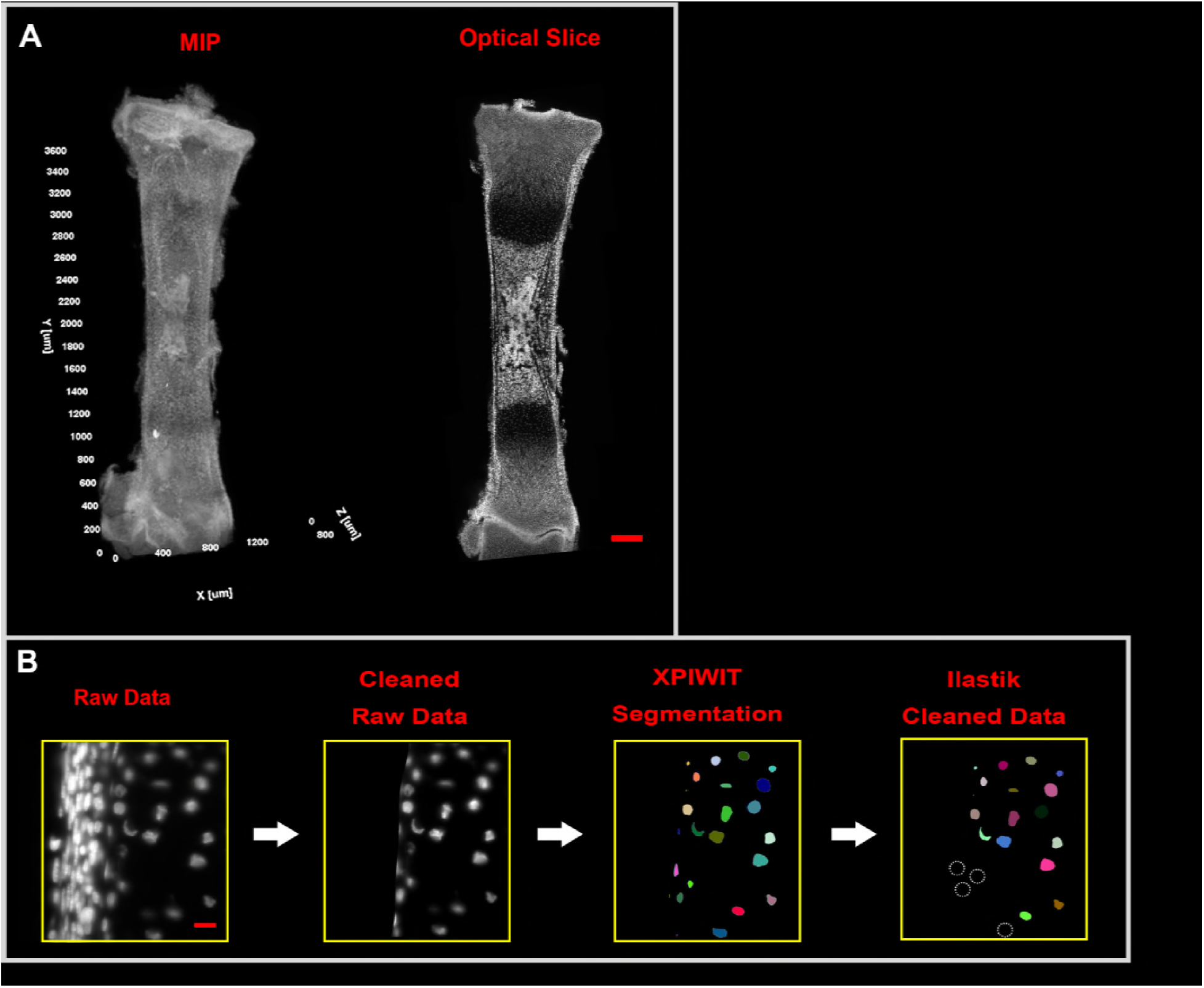
3D imaging and segmentation of growth plate nuclei. **A)** Maximum intensity projection (MIP) image and longitudinal optical slice of cleared E16.5 tibia with DAPI-stained nuclei. Scale bar, 250 μm. **B)** The segmentation pipeline. Raw images are manually segmented in MicroView to remove non-cartilaginous nuclei. Nuclei from the cleaned raw data are automatically segmented in XPIWIT. The resulting segmented nuclei undergo object classification in Ilastik, where incorrect segmentations are removed. Scale bar, 20 μm.

The next challenge was to conduct accurate and time-efficient 3D segmentation of nuclei from the light-sheet images. For that, we combined three open-source platforms to perform semi-automatic segmentations (SAS). Nuclei from surrounding tissues were manually excluded by masking using Microview 2.1.2 (GE Healthcare). Next, the cleaned raw images were automatically segmented using XPIWIT software (32, 33). Then, an object classifier to identify and remove incorrect segmentations was trained using Ilastik (34). 30 segmented images from various regions within the growth plate were used in the training dataset. We were then able to classify hundreds of remaining images blindly by batch processing, thereby greatly speeding up the cleaning stage (Fig 2B).

One of the consequences of imaging a whole tissue was that the border of nuclei residing in the center of the tissue could appear fuzzy due to lower signal-to-noise ratios (Fig S1.), which could obviously influence the accuracy of subsequent analysis. To overcome this potential problem, we performed a benchmark analysis comparing segmentations from two different imaging modalities, namely confocal and light-sheet microscopy. To establish the “ground truth” (GT), nuclei from the four zones of the growth plate, namely the resting (RZ), proliferating (PZ), prehypertrophic (PHZ), and hypertrophic zone (HZ), were manually segmented from confocal images in Microview. Volume and surface area were extracted and corresponding histograms were compared between manual and semi-automatic segmentations using chi-square distance. Results showed that nuclear segmentations by SAS were indistinguishable from manual GT segmentations (p > 0.05), demonstrating the validity of our measurements (Fig 3A). Since the segmentation was performed to the same quality on four different growth plates from two separate litters (Fig 3B), we could use the segmentation error as a tool to estimate the total number of growth plate nuclei at this developmental stage (see Methods). We estimated the number of nuclei to be 178,825±35,582 in the proximal growth plate and 90,460±16,448 in the distal growth plate (Fig 3C).

**Figure 3.**
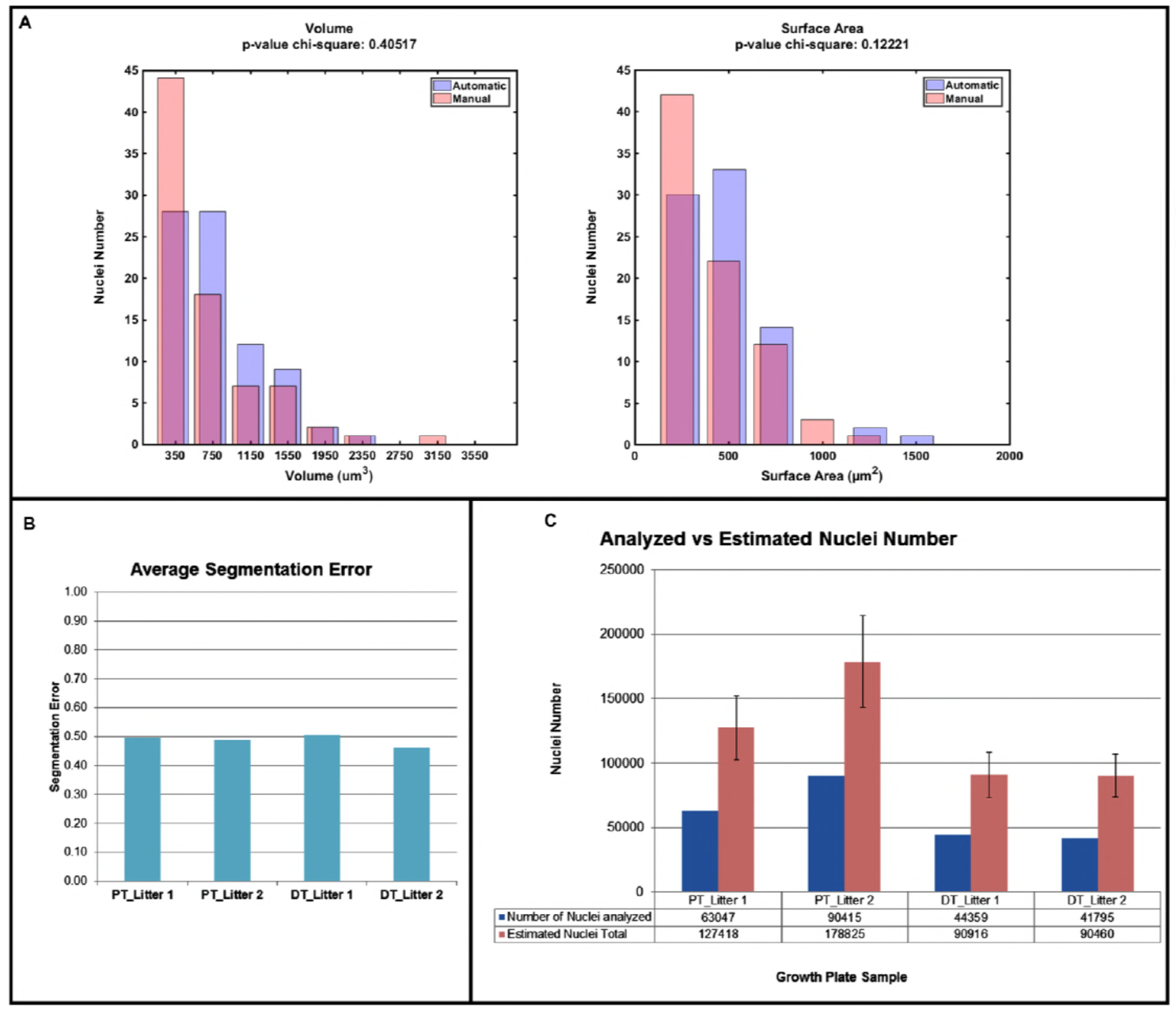
Segmentation validation. **A)** Nuclei from different growth plate zones were imaged by confocal microscopy, manually segmented, and their volumes and surface areas were compared to those of nuclei that were segmented semi-automatically from light sheet images (n=80) using chi-square distance test, resulting in non-significant differences (volume, p = 0.40517; surface area, p = 0.12221). **B)** The average segmentation error was calculated for regions in different growth plate samples by calculating the ratio of correctly segmented nuclei out of the total nuclei in a z-stack (proximal tibia (PT), n = 6; distal tibia (DT), n = 4), resulting in similar error levels across samples. **C)** Bar graph shows the estimated total number of nuclei in each growth plate, based on the number of nuclei analyzed in proximal and distal tibial growth plates from two different litters and the average segmentation error. Error bars represent standard deviations between sample litters.

### Morphometric and spatial analysis of growth plate nuclei

Having segmented successfully nuclei from each growth plate, we proceeded to conduct morphometric analysis on these nuclei, which included calculating the average volume (Fig 4A), surface area (Fig 4B), sphericity (Fig 4C), and spatial orientation (Fig 4F). To understand the spatial distribution between different nuclei, we calculated nuclear density (Fig 4D) and occupation (Fig 4E). Finally, to explore the relationship between nuclear morphology and its location in the growth plate, we stitched the data together and generated 3D morphology maps. For that, nuclei were mapped back to their original locations in the growth plate by using the centroid of each segmented nucleus and the stage coordinates of each image. To analyze data pertaining to tens of thousands of nuclei, we reduced the data dimensionality by splitting the 3D images into a regular grid composed of 75-μm^3^ cubes.

**Figure 4.**
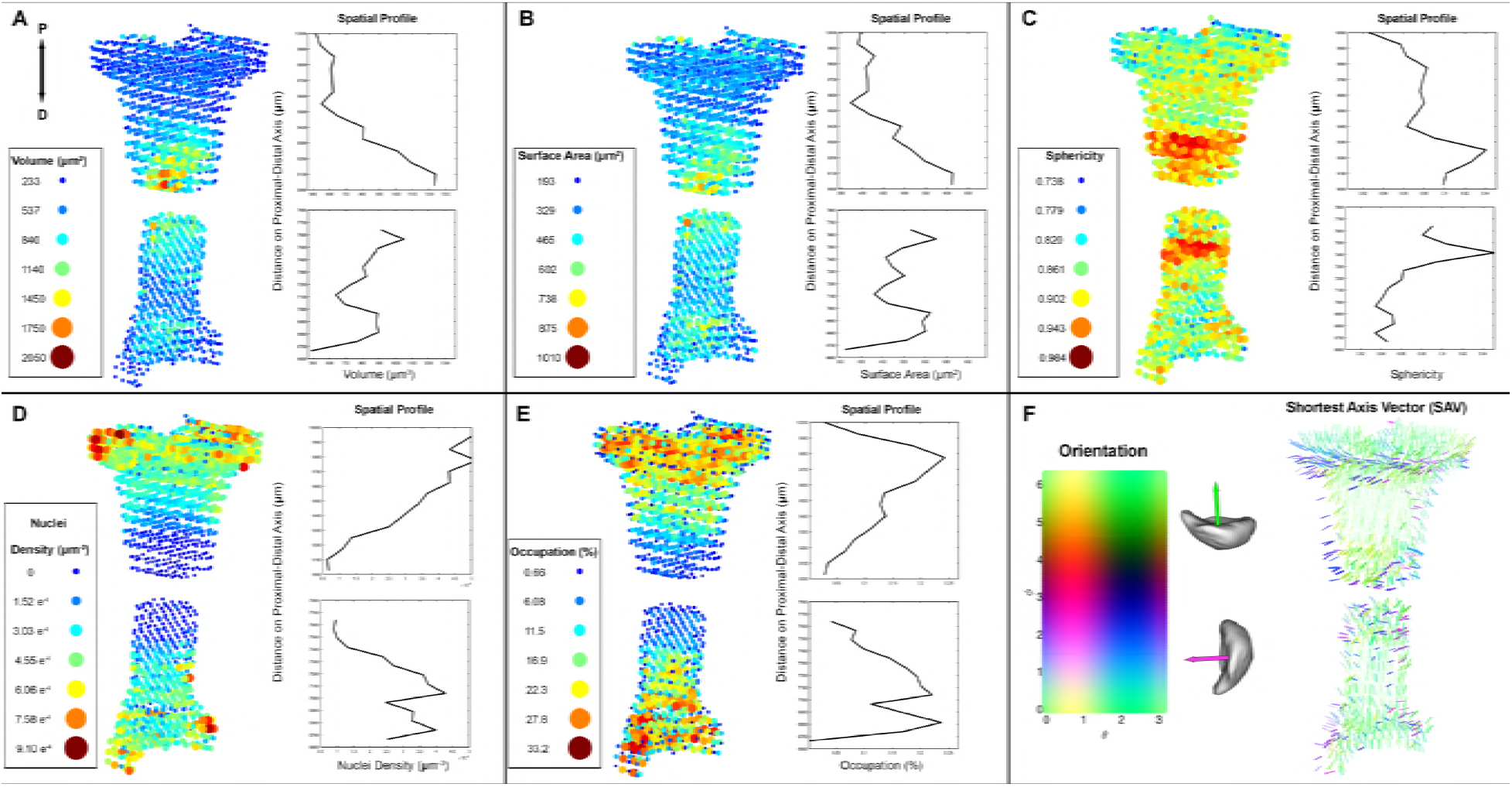
Morphometric and spatial analyses of growth plate nuclei. Segmented nuclei were mapped back to their anatomical position and each growth plate was divided into a regular grid composed of 75 μm cubes. Then, volume (A), surface area (B), sphericity (C), nuclear density (D), occupation (E), and orientation of the shortest axis vector (F) were computed. Colored circles represent the measured values per grid cube. Graphs: The gradient of each measured variable along the proximal-distal axis of the growth plate (“spatial profile”) was computed through the center of each growth plate.

In the proximal growth plate, nuclei gradually increased their volume and surface area up to 9-fold as they differentiated. Interestingly, however, nuclei in the distal growth plate did not form a clear gradient. Instead, there were two regions, in between the presumptive RZ and PZ and in the HZ, where nuclei displayed increased volume and surface area compared to the rest of the tissue. Moreover, maximum volume and surface area were on average 17% and 10% less than in the proximal side, respectively. Relying on previously published data on chondrocyte volume during differentiation (35, 36), we found that nuclear to cell (N/C) volume ratios increased from 0.18 in the RZ to 0.39 in the HZ, suggesting that these ratios are not constant during chondrocyte differentiation in the growth plate. Nuclei located on the edges of the growth plate along the medial-lateral axis consistently had lower volumes and surface areas. Previous studies showed that cells in the RZ and HZ exhibit equally high sphericities (35, 36). By contrast, the highest nuclear sphericities were found in the PHZ and HZ (Fig 4C).

Next, we evaluated nuclear density in the growth plate (Fig 4D). In the proximal growth plate there was a constant gradient along the proximodistal (P-D) axis with highest densities in the RZ and lowest densities in the HZ and along the edges of the growth plate. In the distal growth plate the gradient had two separate phases, a constant increase and a constant decrease along the P-D axis. To study the ratio between nuclear volume and total tissue volume, we calculated the nuclear occupation percentage (Fig 4E). In correlation with our density findings, areas with high nuclear density such as in the RZ had high nuclear occupation and vice versa. To investigate whether different growth plate zones are characterized by unique nuclear morphology features, we performed automated classification of individual nuclei (see Methods) (Fig 5A). Interestingly, the spatial distribution of the classified zones recapitulated the normal organization in the growth plate with 74% accuracy, suggesting that the information encoded in nuclear morphology is sufficient to define a growth plate zone (Fig 5B).

**Figure 5.**
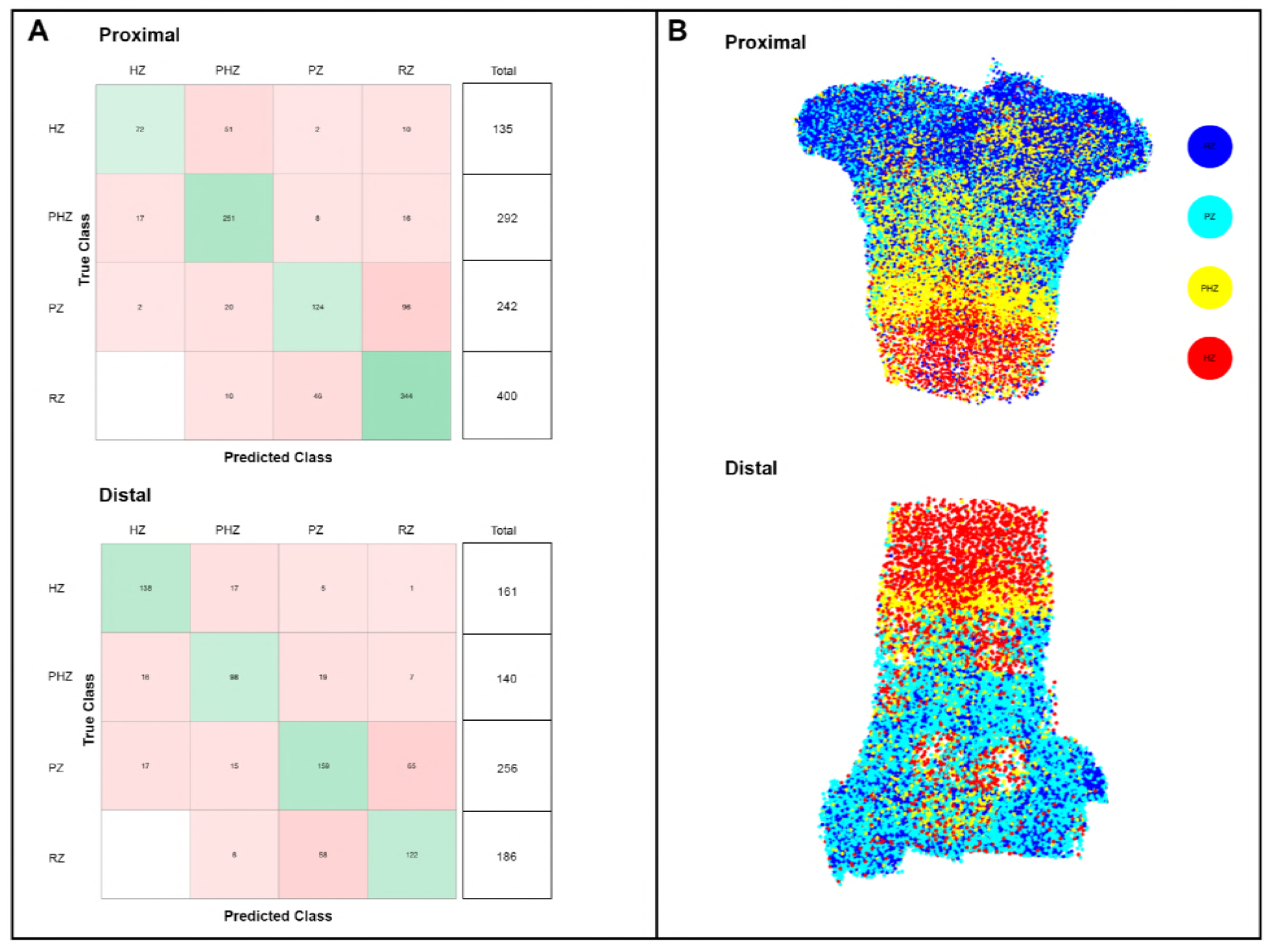
Automatic classification of nuclei into growth plate zones. Manually labeled nuclei from RZ, PZ, PHZ, and HZ were used to train a series of classification models based on their volume, surface area, sphericity, and density. The proximal growth plate was classified with a Fine Gaussian SVM with 74% accuracy, and the distal growth plate was classified by Boosted Tree with an accuracy of 69.5%. The confusion matrix (A) for each growth plate describes the training success. Then the trained classifiers predicted which zone each nucleus belonged to. A 400 μm longitudinal section through the growth plates shows that nuclear morphology is enough to identify clusters representing the four zones (B).

Previous studies showed that cells in the proliferative zone orient themselves into columns that are parallel to the long axis of the bone, facilitating bone elongation (37-40). Using our segmented data, we directly quantified the orientation of the three major axes of each nucleus using principle component analysis (PCA), allowing us to visualize the spatial distribution of nuclear orientations in the growth plate (Fig 4F). In accordance with the current knowledge of cell orientation, we found that nuclei in the PZ also oriented their shortest axis (PC3) along the long axis of the bone with little variability. This trait was conserved in other regions of the growth plate, yet with higher variability (Fig S2). By contrast, a high degree of variability in nuclear orientation direction was observed in the HZ and some areas in the RZ. Finally, nuclei located on the edge of the growth plate oriented orthogonally to the surface of the growth plate.

## Discussion

Data that describes tissue architecture using only one scale, namely tissue or cells, provides limited information. Without the ability to visualize subcellular morphology and location, it is impossible to identify resulting pattern formation within a tissue. This multiscale approach has recently been utilized in studying epithelial tissue morphogenesis in *Drosophila* wings (41) and spheroid spatial heterogeneity (42). In these works, multiscale analysis revealed a role for convergent extension in shaping wing veins, and showed that in breast carcinoma spheroids, there are differences in cell densities between different layers, which should be considered and further explored when modeling this type of cancer. Similar to these recent studies, the method we describe here was developed to gain a system-level understanding of tissue architecture through multiscale analyses of nuclear morphology in the growth plate. By combining tissue clearing and LSFM with high-throughput nuclear segmentation, we were able to implement our newly developed algorithms to create high-resolution 3D tissue maps and extract patterns of nuclear morphology and spatial orientation within the large and complex embryonic growth plate.

For high-throughput extraction of nuclear morphology, our data needed to be acquired at high resolution. Imaging at high resolution creates technical challenges, such as longer imaging time and larger resulting datasets. By acquiring z-stacks consisting only of relevant ROIs, imaging speed was increased and data size was reduced. Even so, our datasets were 500-800 GB in size per growth plate. To overcome this problem, we performed down-sampling of each image. As the voxel dimensions in the raw image (0.1 μm) are ~60 times smaller than the average nucleus diameter (6 μm), isotropic down-sampling by a factor of 2 can be done with negligible loss of morphological accuracy. Several rounds of down-sampling performed before and after segmentation resulted in 80% reduction in data size, greatly speeding up the subsequent preprocessing and segmentation steps.

In recent years, many segmentation tools have become openly available. For example, Nuclear Morphometric Analysis (NMA) (Filippi-Chiela, Oliveira et al. 2012) automatically segments cells and nuclei from 2D tissue culture samples, and NucleusJ (Poulet, Arganda-Carreras et al. 2015) segments nuclei semi-automatically from small 3D image stacks. Yet, neither of these tools is suitable for large 3D datasets, because the first is designed for 2D sparse samples, and the other is computationally slow (Poulet, Arganda-Carreras et al. 2015). Notably, Adaptive Generic Iterative Thresholding Algorithm (AGITA) (Gul-Mohammed, Arganda-Carreras et al. 2014) integrates machine learning to increase speed and accuracy of the segmentation. However, the analysis is paired with the segmentation output and does not allow for quantification of volumes or surface areas.

To overcome the problem of speed and accuracy, we performed automatic segmentation in XPIWIT followed by automatic classification of correct and incorrect segmentations using a trained classifier in Ilastik to quickly extract the 3D morphology of tens of thousands of nuclei per dataset. Although manually training the classifier initially took between one and two weeks for one growth plate, this same training was applied on other untrained growth plates, resulting in classification of the remaining growth plates in one day. As the entire segmentation process is semi-automatic, the user still needs to do some work, such as removing regions in the image that do not belong to the growth plate and tuning segmentation parameters if necessary after evaluating the segmentation output. Additional speed and automation could be gained by initiating these processes directly through computer commands.

Accurate segmentation of thousands of growth plate nuclei allowed us to investigate the relationship between patterns of nuclear morphology and the growth plate. It has been shown in previous studies that there is a correlation between cell and nuclear morphology (43-45). In *Drosophila* egg chambers, nuclear volume strongly correlates with cell volume (46). Additionally, in growth plate explants, shape factors such as width/height ratio as well as volume are correlated between cells and nuclei (47). Previous studies showed that chondrocyte volume increases by up to 6-fold while they move from the resting zone to the hypertrophic zone (35, 36). In agreement with these studies, nuclear volume increased as the cells moved along the growth plate; yet, surprisingly, nuclear volume increased up to 9-fold. Numerous works have shown that nucleus size is tightly linked to cell size. In both growing yeast cells and *Xenopus laevis*, the nucleus to cell (N/C) ratio is constant across cells differing in size (48-51). Interestingly, our measurements for 50% of all growth plate nuclei revealed distinct N/C ratios in different zones. This finding may suggest that in the growth plate the mechanism that preserves the N/C ratio is variable between the different zones, or that it is not functional during chondrocyte differentiation.

Looking at the orientation maps, we were able to identify the presumptive proliferative zone in both growth plates, where cells stereotypically form columns parallel to the long axis of the growth plate (38). In fact, we found that most nuclei in all zones aligned their shortest axis parallel to the main axis of the growth plate, although the degree of variability was noticeably higher in regions other than the proliferative zone. These findings are in line with previous studies of chondrocyte orientation where it was shown that cells in the proliferative zone orient their longest axis perpendicular to the long axis of the growth plate (52-54).

Although the two growth plates of the same bone appear to have similar structures, we could identify some morphological variation between them. For instance, while similar patterns of nuclear density, occupation, and orientation were observed, there were differences in volume, surface area, and sphericity. While there was a constant gradient of volume and surface area through the center of the proximal growth plate, the distal end displayed a gradient with a constant increase and constant decrease along with reduced volume and surface area relative to the proximal side. Past studies have outlined morphological and functional differences between the proximal and distal growth plates of different long bones, particularly in overall size and shape and in growth rate (55-59). The different growth rates between proximal and distal growth plates were recently shown to serve a functional purpose, allowing isometric scaling of symmetry-breaking elements during long bone development (60). Additionally, ATAC-seq on proximal and distal femoral growth plates revealed differences in chromatin accessibility (61). These findings support the notion that in addition to the generic mechanisms that regulate all growth plates, there are also unique features that are regulated separately in each growth plate. As nuclear morphology is tightly linked to chromatin accessibility (62) and gene regulation (63), it is plausible that the changes in nuclear morphology observed in different areas of the growth plate reflect changes in gene expression. Thus, our method may provide a new avenue to study the relationship between tissue structure and gene expression in any tissue through the use of 3D mapping of nuclear morphology.

## Methods

### Animals

For genetic labeling of chondrocyte lineage, *mTmG:Col2a1-Cre* mice were used in all analyses. To create *mTmG:Col2a1-Cre* mice, animals homozygous for *mTmg* (Jackson Laboratories) were crossed with *Col2a1-Cre* mice (Jackson Laboratories). Embryos were dissected in cold phosphate buffered saline (PBS), fixed for 3 hours at 4°C in 4% paraformaldehyde (PFA), washed in PBS and stored at 4°C in 0.5 M EDTA (pH 8.0, Avantor Performance Materials) with 0.01% sodium azide (Sigma). In all timed pregnancies, the plug date was defined as E0.5. For harvesting of embryos, timed-pregnant female mice were sacrificed by carbon dioxide (CO2) exposure. Embryos were sacrificed by decapitation with surgical scissors. Tails were visualized with fluorescent binoculars for genotyping when possible; alternatively, tail genomic DNA was used for genotyping by PCR. All animal experiments were pre-approved by the Institutional Animal Care and Use Committee (IACUC) of the Weizmann Institute.

### Tissue clearing

E16.5 tibias from *mTmG:Col2a1-Cre* mice were cleared using the PACT-deCAL technique (7, 8). Shortly, decalcified samples were washed in PBS, then embedded into a hydrogel of 4% (wt/vol) acrylamide in 1x PBS with 0.25% thermal initiator 2,2’-azobis[2-(2-imidazolin-2-yl)propane]dihydrochloride (Wako, cat. No. VA-044). The hydrogel was allowed to polymerize at 37°C for 3 hours. The sample was removed from the hydrogel, washed in PBS, and moved to 10% SDS with 0.01% sodium azide, shaking at 37°C for 4 days, changing the SDS solution each day. Samples were washed four times with 1x PBST (PBS + 0.1% Triton X-100 + 0.01% sodium azide) at room temperature (RT) over the course of a day. To label nuclei, samples were submerged in 8 μg/ml DAPI in 1x PBST gently shaking overnight at RT. Samples were washed again with four changes of 1x PBST, and the refractive index (RI) of the sample was brought to 1.45 by submersion in a refractive index matching solution (RIMS) consisting of Histodenz (Sigma) and phosphate buffer, shaking gently at RT for 2-3 days. Samples were embedded in 1% low gelling agarose (Sigma) in PBS, in a glass capillary (BRAND, Germany). Embedded samples were submerged in RIMS and protected from light at RT until imaging.

### Light-sheet microscopy

The cleared samples were imaged using a light sheet Z1 microscope (Zeiss Ltd.) equipped with 2 sCMOS cameras PCO-Edge, 10X illumination objectives (LSFM clearing 10X/0.2) and Clr Plan-Neofluar 20X/1,0 Corr nd=1.45 and detection objective dedicated for cleared samples in water-based solution of final RI of 1.45. A low resolution image of the entire tibia was taken with the 20x Clarity lens at a zoom of 0.36. To acquire higher resolution images of the proximal and distal growth plates, multiview imaging was done with the same lens at a zoom of 2.5 resulting in x,y,z voxel sizes of 0.091 μm, 0.091 μm, 0.387 μm. The DAPI and Col2Cre-mGFP channels were acquired with Ex’ 405 nm Em’BP 420-470 and Ex’ 488 nm Em’ BP 505-545 lasers at 2.4% and 3% laser powers, respectively. Light-sheet fusion of images was done if necessary in Zen software (Zeiss). Stitching of low resolution images was done in Arivis Vision4d software (Arivis).

### Image segmentation

Prior to segmentation, images were down-sampled in the X,Y direction using the Downsample plugin in Fiji (64) resulting in x,y,z voxel size of 0.137 μm, 0.137 μm, 0.387 μm. Non-cartilaginous nuclei were manually removed using Microview 2.1.2 (GE Healthcare). Shortly, contours were drawn around non-cartilaginous nuclei, and the resulting ROI was reassigned a pixel intensity value of 0. To automatically segment fluorescently-labeled nuclei in the 3D images, we performed a two-step procedure. In the first step, seed points that were roughly located in the center of the nuclei were detected using a Laplacian-of-Gaussian-based (LoG) approach as described in (33). In brief, the 3D input images were filtered with a LoG filter using a standard deviation that was empirically tuned to the radius of the objects of interest. We used standard deviations of *σ* = 16, *σ* = 18 and *σ* = 45 for RZ/PZ, PHZ, and HZ nuclei, respectively. Subsequently, local intensity maxima were extracted from the LoG-filtered image and reported as potential nuclei centers. For each potential seed point, we compute the average intensity in a 3×3×3 voxel-wide cube surrounding the centroid. In order to minimize the number of false positive detections, only seed points with an average intensity larger than the global average intensity of the entire LoG-filtered image were kept for further processing. In a final step, we used a seeded watershed algorithm to perform the segmentation of the nuclei in a Gaussian-smoothed 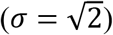 version of the intensity-inverted raw input image. The detected seed points were used to initialize the seeded watershed algorithm and we artificially added a background seed located at the border of each image snippet to separate the centered nucleus from the surrounding background.

The segmentation was performed separately for each nucleus and in parallel, *i.e*., small 3D image patches surrounding the seed points were processed concurrently using multiple cores of the CPU. Segmentation results of the individually processed patches were then combined to form a full-resolution segmentation image containing the final result with a unique integer label for each of the nuclei that was used for further quantification and morphological analyses. All image analysis pipelines were implemented using the open-source software tool XPIWIT (32) and executed on a Windows Server 2012 R2 64-bit workstation with 2 Intel(R) Xeon(R)CPU E5-2690 v3 processors, 256 GB RAM, 24 cores and 48 logical processors.

Segmented images were down-sampled again in the X,Y direction using the Downsample plugin in Fiji (64) resulting in x,y,z voxel sizes of 0.194 μm, 0.194 μm, 0.387 μm. Images were then batch-converted to Hdf5 files using a macro written in Fiji and classified as either “good” or “bad” in Ilastik (34). A total of 37 classified images were used to batch-process the rest of images (n = 240). The resulting classification of the batch-processed images were inspected in Fiji by overlaying the classified image with the raw image and evaluating visually 3D voxel overlap. Finally, voxels belonging to badly segmented nuclei were replaced with a value of 0 in Fiji and images were saved as a 3D tiff stack.

### Nucleus feature extraction

Following image segmentation, we performed an initial data cleaning step that included removal of nuclei that overlapped with the border of the imaged section, as well as nuclei with irregularly small (< 100 μm^3^) or large (> 4000 μm^3^) volumes. To extract morphological features, each nucleus was first converted from a binary volume into a triangle mesh by applying Gaussian smoothing filter (sd = 0.5 pixels) and extracting the iso-surface at the iso-value 0.5. From the triangle mesh of each nucleus, the following features were extracted: surface area, as the sum of the areas of all mesh faces; volume, using the convergence theorem (a.k.a. Gauss’s theorem or Ostrogradsky’s theorem (65)); sphericity, calculated as:

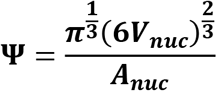

where ***V_nuc_*** is the volume of the nucleus and ***A_nuc_*** is the surface area (66); nucleus orientation, defined as the direction of the third principal component of the point-cloud of vertices of the triangular mesh, which serves as a normal to the largest face of the nucleus’s surface. Feature extraction was done on an Intel^®^ Xeon^®^ CPU E5-1620 v4 @ 3.50GHz, 32.0 GB RAM, Windows 10 64 bit, with MATLAB version 2017b (The MathWorks, Inc., Natick, Massachusetts, USA).

### Machine learning classification

To evaluate how well nuclear morphology could predict to which zone a cell belongs, we developed an automated classification method using MATLAB R2018a’s “Classification Learner toolbox”. We used classification models on manually labeled nuclei based on their volume, surface area, sphericity and density to train a series of classifiers (Table S1). The feature values were first normalized as a score function (i.e., subtraction of the mean value and division by the standard deviation obtained from the entire set of values). The accuracy of each of the classification models was assessed using a 5-fold cross-validation method. The best model for the classification of the nuclei in the proximal growth plate was a Fine Gaussian SVM, with an accuracy of 74%, whereas the best model for the classification of the nuclei in the distal growth plate was Boosted Tree with an accuracy of 69.5%. Confusion matrices describe the details of the prediction. Using the learned classifiers, we made predictions on the unlabeled nuclei.

### Segmentation Validation

To validate segmentation, an E16.5 tibia from a sample littermate was dissected and cleared using PACT-deCAL, stained with DAPI, mounted in RIMS on a glass slide with a shallow well, and covered with a glass coverslip. 75-μm z-stacks from areas in the RZ, PZ, PHZ, and HZ were acquired with x,y,z voxel sizes of 0.1 μm, 0.1 μm, 0.5 μm. Confocal images were manually segmented (n = 80 nuclei) in Microview software and volume and surface area were calculated for each nucleus. Volume and surface area of automatically segmented nuclei (n = 80) from comparable regions in the growth plate were compared to the manual segmentations by a chi-square distance between histograms test (67). We could not reject the hypothesis that the two distributions are drawn from the same underlying distribution (p > 0.05), demonstrating that our semi-automatic segmentations were valid.

### Segmentation error calculation

To calculate the segmentation error from six representative regions in the proximal and five representative regions in the distal growth plate, we calculated the average ratio of analyzed nuclei out of the total nuclei number in each region of 2D optical sections every 23 μm along the Z axis, to avoid counting the same nucleus twice. To count the total nuclei number in each slice, we counted nuclei in the DAPI channel, using a macro written in FIJI. To count the number of analyzed nuclei out of each slice, we used the MATLAB scripts. The resulting calculations were done using the following equations:

Average error = total nuclei / analyzed nuclei
Estimated total nuclei = # analyzed nuclei * (Average error)
STD = std (Average error)
Average segmentation error = 1 / average error per growth plate

### 3D nuclear morphology maps of growth plates

To visualize the spatial distribution of nuclear morphology within each growth plate, data were first stitched back together in MATLAB using the stage coordinates and scaling of each image. To place each voxel in a global coordinate system, we added the coordinates of each stack to every voxel in the stack, thereby reconstructing the entire bone. To highlight large patterns while averaging out small differences between individual nuclei, we chose to represent nuclear features at a coarse-grained level. This representation was computed by first defining a regular spatial grid over the data. Each element of the grid was defined as a 75 x 75 x 75 μm^3^ cube, containing up to 275 nuclei. Within each cube, we averaged the nucleus volume, surface area and sphericity and used the Delaunay tessellation field estimator at a resolution of 2 μm^3^ to compute the nuclear density for each cube (68). In addition, for each cube, we computed the occupation, defined as the sum of nuclear volumes divided by the volume of the cube. Then, using MATLAB’s jet color map, we represented the characteristics of each cube on the grid by drawing spheres whose radii and colors are proportional to the computed values and whose centers correspond to the average position of the nuclei within the bin.

### Nucleus orientation maps and scatter plots

The spatial distribution of nuclear orientations was visualized using a regular spatial grid of the same size as the morphology maps. The orientation of a nucleus is described by the third principal component of a PCA performed on the point cloud corresponding to the segmented shape of the nucleus. When computing the principal component, the sign of the vector is attributed arbitrarily. Before averaging the orientations in a cube, we first constrained the orientations to be in the same hemisphere to avoid artificial bias. This was performed by considering all the orientations and their opposites, leading to 2N vectors, where N is the number of nuclei in the cube. We then performed a PCA on the set of orientation vectors to determine the main direction. The N orientation vectors corresponding to the hemisphere representing the main direction were extracted by considering only the ones having a positive dot product with the main direction. The main orientation is computed over the N selected orientation vectors as a standard average over the direction cosines (69). The spherical variance is computed as the dispersion around the mean direction (69).

To compare among growth plates, we registered the stitched data beforehand. The registration was performed in two steps using MATLAB’s rigid registration. First, each growth plate was registered to an artificially generated set of points mimicking the global shape of a growth plate, with the P-D axis aligned with the X axis. Then, each growth plate was registered to one of the samples. For each cube a line was drawn oriented and color-coded according to the average orientation, and whose thickness was proportional to the spherical variance. Finally, to summarize the distribution of orientations in a 2D graph, we used the fact that each of the mean orientation vectors obtained for each of the bin of the regular grid were encoded using spherical coordinates (*θ* and *ϕ*, for polar and azimuthal angles respectively). *θ* varies from 0 to *π*, while *ϕ* varies from —*π* to *π*, enabling the encoding of any orientation in 3D.

### Spatial profiles of morphology features

To visualize quantitatively the spatial profile of each of the computed features and identify gradient along the growth plate, we defined an irregular grid as following: The entire growth plate was divided into 75 μm-thick slices along the P-D axis, and into three equally spaced slices along each of the two remaining axis. This division resulted in 9 rods organized in a 3 x 3 formation and subdivided at 75 μm intervals along the P-D axis. The spatial profiles were computed by averaging the individual cell features in each of the resulting cubes in the most central rod only.

## Acknowledgements

We thank Nitzan Konstantin for editorial assistance and members of the Zelzer lab for their advice and encouragement throughout this project. We thank the de Picciotto-Lesser Cell Observatory in memory of Wolf and Ruth Lesser, Weizmann Institute of Science, for providing LSFM infrastructure, Rada Massarwa and Jacob Hanna from the Department of Molecular Genetics, Weizmann Institute, for providing *mTmg* mice, Ofra Golani from the MICC Cell Observatory and Kiril Kogan from the Bioinformatics Unit, Weizmann Institute, for support and discussions regarding segmentation analysis, and Tali Wiesel from the Graphic Design Department at the Weizmann Institute of Science for her help with graphics.

This study was supported by grants from the National Institutes of Health (NIH, #R01 AR055580), European Research Council (ERC, #310098), the Israel Science Foundation (ISF, #345/16), the David and Fela Shapell Family Center for Genetic Disorders, and the Estate of Bernard Bishin for the WIS-Clalit Program (to E.Z), the WIN program between Princeton University and the Weizmann Institute (to P.V.), and from the German Research Foundation (DFG-MI1315/4-1; to J.S).

---

**Figure S1.**
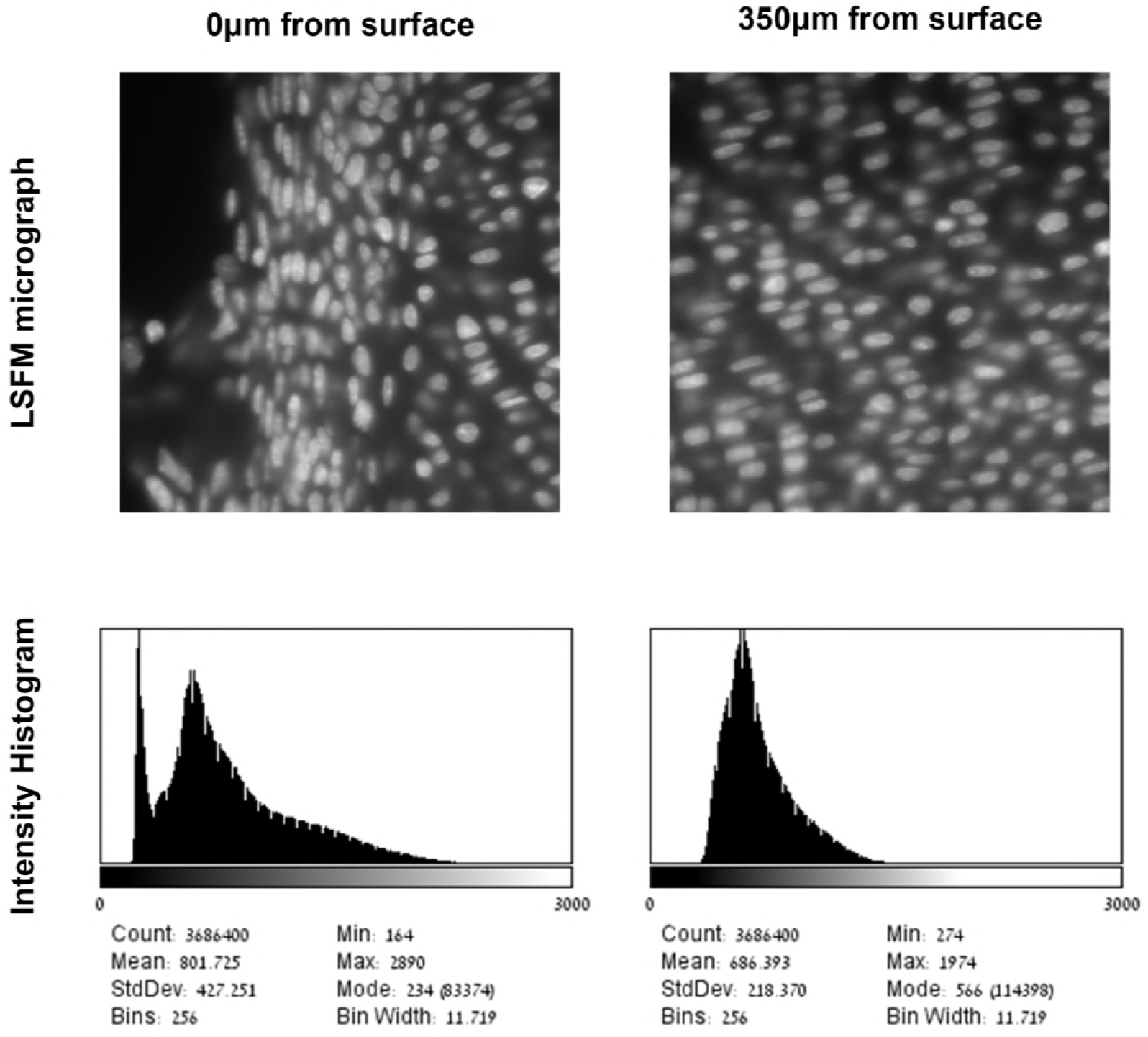
Signal-to-noise ratio is reduced away from the growth plate surface (related to Figure 3). Optical sections from LSFM imaging of cleared growth plates with their accompanying intensity histograms generated in Fiji (below) demonstrate that the signal-to-noise ratio in nuclear images is lower in the center of the tissue than on the edge.

**Figure S2.**
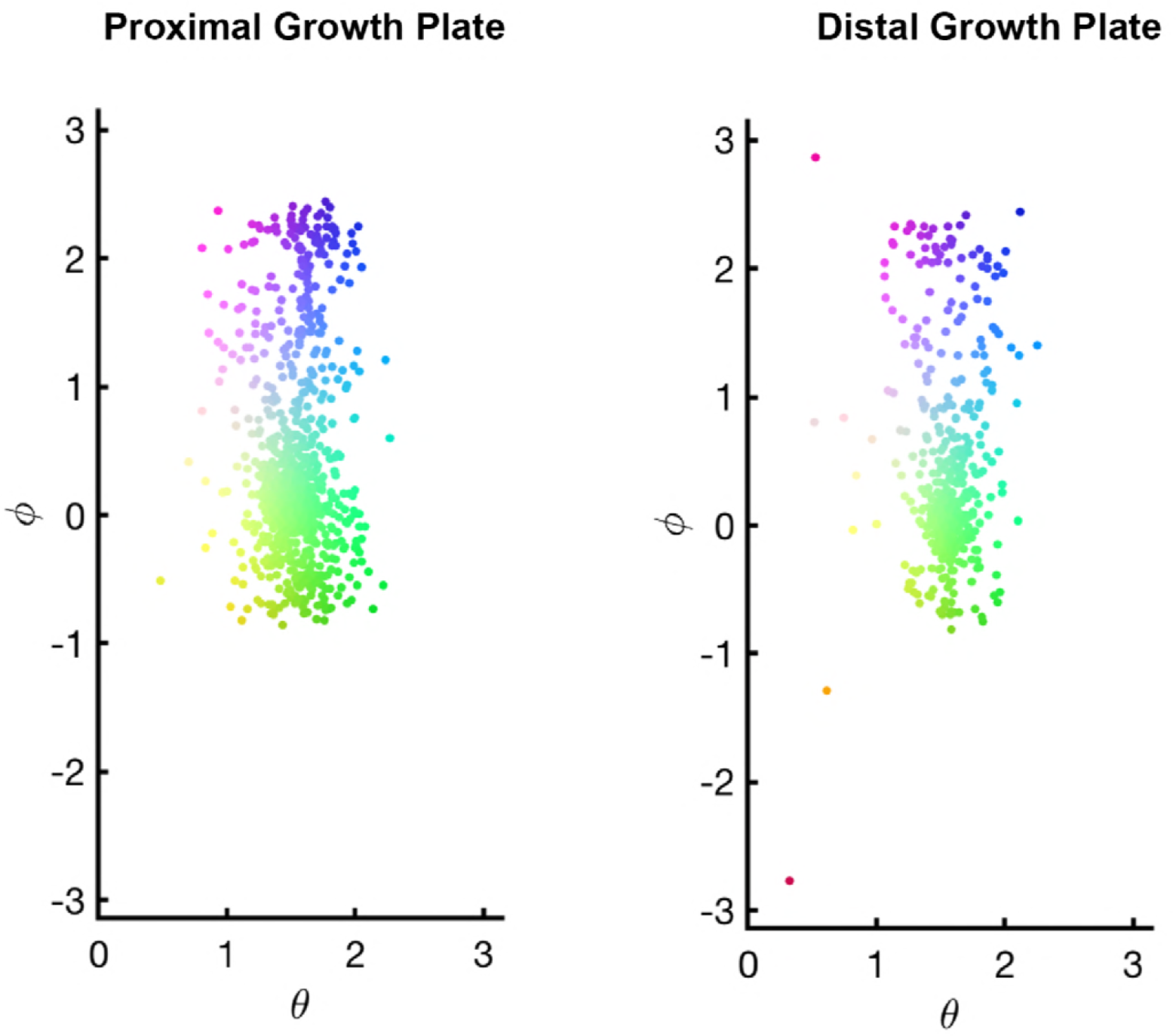
Spatial distribution of nuclear orientation (related to Figure 4). Scatter plots of nuclear orientation defined by polar angle *θ* and azimuthal angle *ϕ* for the proximal and distal growth plates. Colored dots represent distribution of the colored lines in Figure 4F. Large concentration of green dots highlights the existence of a preferential nuclear orientation in both growth plates.

**Table S1.** Proportions of manually labeled nuclei in each growth plate zone (related to Figure 5). Automatically segmented nuclei from the proximal and distal growth plates were labeled as belonging to the HZ, PHZ, PZ, or RZ. The volume, surface area, sphericity, and density of these labeled nuclei were used to train a series of classifiers to predict which zone they belong to.

|  | HZ | PHZ | PZ | RZ | Total number of labeled nuclei | Total number of segmented nuclei |
| --- | --- | --- | --- | --- | --- | --- |
| Proximal | 136 | 295 | 279 | 459 | 1169 | 90415 |
| Distal | 165 | 140 | 304 | 207 | 816 | 41795 |

## References

1. Kaufmann A, Mickoleit M, Weber M, Huisken J. Multilayer mounting enables long-term imaging of zebrafish development in a light sheet microscope. Development. 2012;139(17):3242.

2. Keller PJ, Schmidt AD, Wittbrodt J, Stelzer EHK. Reconstruction of Zebrafish Early Embryonic Development by Scanned Light Sheet Microscopy. Science. 2008;322(5904):1065.

3. Chardes C, Melenec P, Bertrand V, Lenne PF. Setting up a simple light sheet microscope for in toto imaging of C. elegans development. LID - 10.3791/51342[doi]. (1940-087X (Electronic)).

4. Rieckher M, Kyparissidis-Kokkinidis I, Zacharopoulos A, Kourmoulakis G, Tavernarakis N, Ripoll J, et al. A Customized Light Sheet Microscope to Measure Spatio-Temporal Protein Dynamics in Small Model Organisms. PLOS ONE. 2015;10(5):e0127869.

5. Huisken J, Swoger J, Del Bene F, Wittbrodt J, Stelzer EHK. Optical Sectioning Deep Inside Live Embryos by Selective Plane Illumination Microscopy. Science. 2004;305(5686):1007.

6. Richardson DS, Lichtman JW. Clarifying Tissue Clearing. Cell. 2015;162(2):246–57.

7. Treweek JB, Chan KY, Flytzanis NC, Yang B, Deverman BE, Greenbaum A, et al. Whole-body tissue stabilization and selective extractions via tissue-hydrogel hybrids for high-resolution intact circuit mapping and phenotyping. Nat Protocols. 2015;10(11):1860–96.

8. Yang B, Treweek Jennifer B, Kulkarni Rajan P, Deverman Benjamin E, Chen C-K, Lubeck E, et al. Single-Cell Phenotyping within Transparent Intact Tissue through Whole-Body Clearing. Cell. 158(4):945–58.

9. Calve S, Ready A, Huppenbauer C, Main R, Neu CP. Optical Clearing in Dense Connective Tissues to Visualize Cellular Connectivity In Situ. PLOS ONE. 2015;10(1):e0116662.

10. Greenbaum A, Chan KY, Dobreva T, Brown D, Balani DH, Boyce R, et al. Science Translational Medicine. 2017;9(387).

11. Zink D, Fischer AH, Nickerson JA. Nuclear structure in cancer cells. Nature Reviews Cancer. 2004;4:677.

12. Webster M, Witkin KL, Cohen-Fix O. Sizing up the nucleus: nuclear shape, size and nuclear-envelope assembly. Journal of Cell Science. 2009;122(10):1477.

13. Scaffidi P, Misteli T. Lamin A-Dependent Nuclear Defects in Human Aging. Science. 2006;312(5776):1059.

14. Haithcock E, Dayani Y, Neufeld E, Zahand AJ, Feinstein N, Mattout A, et al. Age-related changes of nuclear architecture in *em*Caenorhabditis elegans*em*. Proceedings of the National Academy of Sciences of the United States of America. 2005;102(46):16690.

15. Khatau SB, Kusuma S, Hanjaya-Putra D, Mali P, Cheng L, Lee JSH, et al. The Differential Formation of the LINC-Mediated Perinuclear Actin Cap in Pluripotent and Somatic Cells. PLoS ONE. 2012;7(5):e36689.

16. Skinner BM, Johnson EEP. Nuclear morphologies: their diversity and functional relevance. Chromosoma. 2017;126(2):195–212.

17. Mosser DM, Edwards JP. Exploring the full spectrum of macrophage activation. (1474-1741 (Electronic)).

18. Pagliara S, Franze K, McClain CR, Wylde G, Fisher CL, Franklin RJM, et al. Auxetic nuclei in embryonic stem cells exiting pluripotency. (1476-1122 (Print)).

19. Uhler C, Shivashankar GV. Regulation of genome organization and gene expression by nuclear mechanotransduction. Nature Reviews Molecular Cell Biology. 2017;18:717.

20. Kirby TJ, Lammerding J. Emerging views of the nucleus as a cellular mechanosensor. Nature Cell Biology. 2018;20(4):373–81.

21. Fenelon KD, Hopyan S. Structural components of nuclear integrity with gene regulatory potential. Current Opinion in Cell Biology. 2017;48:63–71.

22. Shivashankar GV. Mechanosignaling to the Cell Nucleus and Gene Regulation. Annual Review of Biophysics. 2011;40(1):361–78.

23. Uhler C, Shivashankar GV. Regulation of genome organization and gene expression by nuclear mechanotransduction. (1471-0080 (Electronic)).

24. Uhler C, Shivashankar GV. Chromosome Intermingling: Mechanical Hotspots for Genome Regulation. Trends in Cell Biology. 27(11):810–9.

25. Allis CD, Jenuwein T. The molecular hallmarks of epigenetic control. (1471-0064 (Electronic)).

26. Gesson K, Rescheneder P, Skoruppa MP, von Haeseler A, Dechat T, Foisner RA-Ohoo. A-type lamins bind both hetero-and euchromatin, the latter being regulated by lamina-associated polypeptide 2 alpha. (1549-5469 (Electronic)).

27. Mackie EJ, Ahmed Ya Fau - Tatarczuch L, Tatarczuch L Fau - Chen KS, Chen Ks Fau - Mirams M, Mirams M. Endochondral ossification: how cartilage is converted into bone in the developing skeleton. (1357-2725 (Print)).

28. Kronenberg HM. Developmental regulation of the growth plate. (0028-0836 (Print)).

29. Cooper KL, Oh S, Sung Y, Dasari RR, Kirschner MW, Tabin CJ. Multiple Phases of Chondrocyte Enlargement Underlie Differences in Skeletal Proportions. Nature. 2013;495(7441):375–8.

30. Abad V, Meyers JL, Weise M, Gafni RI, Barnes KM, Nilsson O, et al. The Role of the Resting Zone in Growth Plate Chondrogenesis. Endocrinology. 2002;143(5):1851–7.

31. Cancedda R, Cancedda FD, Castagnola P. Chondrocyte Differentiation. In: Kwang WJ, Jonathan J, editors. International Review of Cytology. Volume 159: Academic Press; 1995. p. 265–358.

32. Bartschat A, Hübner E, Reischl M, Mikut R, Stegmaier J. XPIWIT—an XML pipeline wrapper for the Insight Toolkit. Bioinformatics. 2015;32(2):315–7.

33. Stegmaier J, Otte JC, Kobitski A, Bartschat A, Garcia A, Nienhaus GU, et al. Fast Segmentation of Stained Nuclei in Terabyte-Scale, Time Resolved 3D Microscopy Image Stacks. PLOS ONE. 2014;9(2):e90036.

34. Sommer C, Straehle C, Köthe U, Hamprecht FA, editors. Ilastik: Interactive learning and segmentation toolkit. 2011 IEEE International Symposium on Biomedical Imaging: From Nano to Macro; 2011 March 30 2011-April 2 2011.

35. Amini S, Veilleux D, Villemure I. Tissue and cellular morphological changes in growth plate explants under compression. Journal of Biomechanics. 2010;43(13):2582–8.

36. Amini S, Veilleux D Fau - Villemure I, Villemure I. Three-dimensional in situ zonal morphology of viable growth plate chondrocytes: a confocal microscopy study. (1554-527X (Electronic)).

37. Williams RM, Zipfel Wr Fau - Tinsley ML, Tinsley Ml Fau - Farnum CE, Farnum CE. Solute transport in growth plate cartilage: in vitro and in vivo. (0006-3495 (Print)).

38. Romereim SM, Conoan NH, Chen B, Dudley AT. A dynamic cell adhesion surface regulates tissue architecture in growth plate cartilage. Development (Cambridge, England). 2014;141(10):2085–95.

39. Kuss P, Kraft K, Stumm J, Ibrahim D, Vallecillo-Garcia P, Mundlos S, et al. Regulation of cell polarity in the cartilage growth plate and perichondrium of metacarpal elements by HOXD13 and WNT5A. Developmental Biology. 2014;385(1):83–93.

40. de Andrea CE, Wiweger M Fau - Prins F, Prins F Fau - Bovee JVMG, Bovee Jv Fau - Romeo S, Romeo S Fau - Hogendoorn PCW, Hogendoorn PC. Primary cilia organization reflects polarity in the growth plate and implies loss of polarity and mosaicism in osteochondroma. (1530-0307 (Electronic)).

41. Etournay R, Merkel M, Popović M, Brandl H, Dye NA, Aigouy B, et al. TissueMiner: A multiscale analysis toolkit to quantify how cellular processes create tissue dynamics. eLife. 2016;5:e14334.

42. Schmitz A, Fischer SC, Mattheyer C, Pampaloni F, Stelzer EHK. Multiscale image analysis reveals structural heterogeneity of the cell microenvironment in homotypic spheroids. Scientific Reports. 2017;7:43693.

43. Dapples CC, King RC. The development of the nucleolus of the ovarian nurse cell of Drosophila melanogaster. Zeitschrift für Zellforschung und Mikroskopische Anatomie. 1970;103(1):34–47.

44. Jacob J, Sirlin JL. Cell function in the ovary of drosophila. Chromosoma. 1959;10(1):210–28.

45. Versaevel M, Grevesse T, Gabriele S. Spatial coordination between cell and nuclear shape within micropatterned endothelial cells. Nature Communications. 2012;3:671.

46. Imran Alsous J, Villoutreix P, Berezhkovskii AM, Shvartsman SY. Collective Growth in a Small Cell Network. Current Biology. 2017;27(17):2670–6.e4.

47. Guilak F. Compression-induced changes in the shape and volume of the chondrocyte nucleus. (0021-9290 (Print)).

48. Levy DL, Heald R. Nuclear size is regulated by importin alpha and Ntf2 in Xenopus. (1097-4172 (Electronic)).

49. Jorgensen P, Edgington NP, Schneider BL, Rupeš I, Tyers M, Futcher B. The Size of the Nucleus Increases as Yeast Cells Grow. Molecular Biology of the Cell. 2007;18(9):3523–32.

50. Hara Y, Merten Christoph A. Dynein-Based Accumulation of Membranes Regulates Nuclear Expansion in Xenopus laevis Egg Extracts. Developmental Cell. 2015;33(5):562–75.

51. Neumann FR, Nurse P. Nuclear size control in fission yeast. The Journal of Cell Biology. 2007;179(4):593.

52. Dodds GS. Row formation and other types of arrangement of cartilage cells in endochondral ossification. The Anatomical Record. 1930;46(4):385–99.

53. Ascenzi M-G, Blanco C, Drayer I, Kim H, Wilson R, Retting KN, et al. Effect of localization, length and orientation of chondrocytic primary cilium on murine growth plate organization. Journal of theoretical biology. 2011;285(1):147–55.

54. Li Y, Dudley AT. Noncanonical frizzled signaling regulates cell polarity of growth plate chondrocytes. Development. 2009;136(7):1083.

55. Wilsman NJ, Bernardini ES, Leiferman E, Noonan K, Farnum CE. Age and Pattern of the Onset of Differential Growth Among Growth Plates in Rats. Journal of orthopaedic research : official publication of the Orthopaedic Research Society. 2008;26(11):1457–65.

56. Breur GJ, Vanenkevort BA, Farnum CE, Wilsman NJ. Linear relationship between the volume of hypertrophic chondrocytes and the rate of longitudinal bone growth in growth plates. Journal of Orthopaedic Research. 1991;9(3):348–59.

57. Church Le Fau - Johnson LC, Johnson LC. GROWTH OF LONG BONES IN THE CHICKEN. RATES OF GROWTH IN LENGTH AND DIAMETER OF THE HUMERUS, TIBIA, AND METATARSUS. (0002-9106 (Print)).

58. Digby KH. The Measurement of Diaphysial Growth in Proximal and Distal Directions. Journal of Anatomy and Physiology. 1916;50(Pt 2):187–8.

59. Payton CG. The Growth in Length of the Long Bones in the Madder-fed Pig. Journal of Anatomy. 1932;66(Pt 3):414–25.

60. Stern T, Aviram R, Rot C, Galili T, Sharir A, Kalish Achrai N, et al. Isometric Scaling in Developing Long Bones Is Achieved by an Optimal Epiphyseal Growth Balance. PLOS Biology. 2015;13(8):e1002212.

61. Guo M, Liu Z, Willen J, Shaw CP, Richard D, Jagoda E, et al. Epigenetic profiling of growth plate chondrocytes sheds insight into regulatory genetic variation influencing height. eLife. 2017;6:e29329.

62. Ramdas NM, Shivashankar GV. Cytoskeletal Control of Nuclear Morphology and Chromatin Organization. Journal of Molecular Biology. 2015;427(3):695–706.

63. Thomas CH, Collier JH, Sfeir CS, Healy KE. Engineering gene expression and protein synthesis by modulation of nuclear shape. Proceedings of the National Academy of Sciences of the United States of America. 2002;99(4):1972–7.

64. Schindelin J, Arganda-Carreras I, Frise E, Kaynig V, Longair M, Pietzsch T, et al. Fiji: an open-source platform for biological-image analysis. Nat Meth. 2012;9(7):676–82.

65. Gauß CF. Theoria attractionis corporum sphaeroidicorum ellipticorum homogeneorum methodo nova tractata1813.

66. Wadell H. Volume, Shape, and Roundness of Quartz Particles: University of Chicago, Department of Geology.; 1932.

67. Porter FC. Testing Consistency of Two Histograms. ArXiv e-prints [Internet]. 2008 April 1, 2008; 0804. Available from: http://adsabs.harvard.edu/abs/2008arXiv0804.0380P.

68. Schaap WE. DTFE: the Delaunay Tessellation Field Estimator s.n.: University of Groningen; 2007.

69. Mardia KV. Statistics of directional data: Academic press; 2014.

